# Is high-frequency activity evidence of an anterior temporal lobe network or micro-saccades?

**DOI:** 10.1101/2023.01.09.523285

**Authors:** George C. O’Neill, Stephanie Mellor, Robert A. Seymour, Nicholas Alexander, Tim M. Tierney, Ryan C. Timms, Eleanor A. Maguire, Gareth R. Barnes

## Abstract

There is renewed interest in electrical activity that extends beyond the typical electrophysiological 100 Hz bandwidth. This activity, often in the anterior temporal lobe, has been attributed to processes ranging from memory consolidation to epileptiform activity. Here, using an open-access resting state magnetoencephalography (MEG) dataset (n = 89), and a second task-based MEG dataset, we could reliably localise high-frequency power to the temporal lobes across multiple bands up to 300–400 Hz. A functional connectivity analysis of this activity revealed a robust resting state bilateral network between the temporal lobes. However, we also found robust coherence in the 100–200 and 200–300 Hz bands between source reconstructed MEG data and the electrooculography (EOG) localised to within the temporal poles. Additional denoising schemes applied to the data could reduce power localisation to the temporal poles but the topography of the functional network did not drastically alter. Whilst it is clear that this network is biological and robust to established denoising methods, we cannot definitively rule yet on whether this is of neural or myogenic origin.

## 1. Introduction

Typical magnetoencephalography/electroencephalography (MEG/EEG) studies focus on the electromagnetic signals from brain activity in the 0.1–100 Hz bandwidth. Physiologically, ripple phenomena extending up to 200 Hz have been identified within the hippocampus and neighbouring temporal lobe structures (Buzsáki, 1989, 1986). One theory is that these discharges (driven by inter-neurons forcing successive firing of multiple subpopulations of pyramidal neurons) facilitate the transfer of transient memory traces from hippocampus to entorhinal cortex (Ylinen et al., 1995). Most recently, MEG activity linked to memory replay was measured non-invasively in the 120–150 Hz band in the 100 ms post-stimulus (Liu et al., 2019). These replay events were found to coincide with a broadband (>100 Hz) increase in anterior temporal lobe network activity (Higgins et al., 2021). Recent work using a non-invasive low-noise magnetometry system has revealed evidence of measurable fields from healthy endogenous brain activity even up to 1000 Hz (Waterstraat et al., 2021).

In the clinical domain, high-frequency signals have been extensively studied in the context of epilepsy and their role as a useful biomarker of the epileptogenic zone (Staba et al., 2002) both invasively and using MEG (Velmurugan et al., 2019; Xiang et al., 2021). Although such invasive recordings are only available from recordings in implanted patients, there are reports of endogenous high-frequency activity in which there is no clear link to epileptogenic or lesioned cortex. For example, Mari and colleagues (2012) reported ‘continuous’ high frequency phenomena in human medial temporal lobe structures which were not significantly influenced by lesion or seizure onset zone. In a subsequent study, Melani et al. (2013) found that these high frequency phenomena (80-250 Hz) were unrelated to seizure onset zone or lesion site but showed a preferential distribution for the hippocampus and occipital lobe. Most recently, combining data from multiple implantations across two centres, Frauscher et al. (2018) produced a map of physiological ripple rates (in non-lesional tissue) during sleep. They categorized the signals into physiological ripples (80–250 Hz), fast ripples (>250Hz) and high-frequency activity (defined as long-lasting oscillation over 80 Hz). They found 80–250Hz ripple events to be ubiquitous and predominate in eloquent cortex, with sustained high-frequency oscillations (>80Hz) and fast ripples (>250Hz) being much rarer in eloquent and non-eloquent regions. However, in contrast to Melani et al. (2018), no dominant hippocampal sites were identified, perhaps because sites with signs of epileptogenic activity were not analysed. These ripple phenomena were also relatively rare (~1/minute).

However, one important concern is that muscle activity also has a broadband electrical signature (Muthukumaraswamy, 2013). This effect was noted in intracranial EEG (iEEG) recordings at the anterior temporal poles where gamma band contamination from *rectus lateralis* muscle occurs (Jerbi et al., 2009). These findings were supported by Kovach et al. (2011) who were able to show the similarity (in spectral, temporal and cross-trial comparisons) between the intracranial temporal pole electrical measurements and occular myographic recordings. Carl et al. (2012) extended these findings to non-invasive MEG recordings in healthy participants and were able to show a saccadic spike source (occupying the 32-128 Hz band) originating in the rectus muscle, but blurring into temporal pole and orbitofrontal areas. Other muscles implicated in apparent intracranial electrical activity include those implicated in grimacing and mastication (Otsubo et al., 2008).

Given the findings summarised above, here we investigated whether signs of high-frequency endogenous activity were evident in two standard, previously acquired, multi-channel MEG recordings. The first dataset was recorded during resting state (Larson-Prior et al., 2013), and we examined the distribution of electrical power peaks throughout the brain. We also reanalysed another independent, task-based dataset previously recorded at our Centre to ascertain if similar effects were evident (Barry et al., 2019).

## 2. Methods

### 2.1 Data and Experiment

#### 2.1.1 Human Connectome Project Resting State Data

Data from 89 subjects from the Human Connectome Project (HCP; Larson-Prior et al., 2013) who underwent MEG data collection were used for this study. Each subject underwent a battery of tests, but here we focus on the first of three 6-minute eyes open resting state scans that were collected. MEG data were acquired on a whole-head Magnes 3600 scanner (4D Neuroimaging, San Diego, CA, USA) at a sample rate of 2034.51 Hz. Participants were placed supine into the system and instructed to fixate on a cross in the centre of the screen. Three head position indicators (HPIs) were placed on landmarks corresponding to the nasion and preauricular areas to allow for registration of the subjects’ anatomy (derived from their 3T MRI scans) to the MEG sensors. In addition to the MEG data, electrodes were placed to measure electrooculography in both horizontal (HEOG) and vertical (VEOG) directions, as well as electrocardiography (ECG). For control analyses to establish the SNR limit of these source data, we also used the supplied 5-minute empty room recording acquired prior to human data collection. For both resting state and empty room recordings, default synthetic gradiometer compensation matrices supplied at data collection were retained and applied to the data.

#### 2.1.2 Scene Imagination Data

We also analysed a second task-based dataset, which has been explored in previous studies (Barry et al., 2019; O’Neill et al., 2021). 22 native English speakers (14 female, aged 27±7 [mean±SD] years) participated in study that involved generating novel scene imagery. The study was approved by the University College London Research Ethics Committee and all participants gave written informed consent.

On any one trial, participants were asked to either imagine a scene, a single isolated object floating against a white background, or count in threes from a specified number. The stimulus type (either, “scene”, “object” or “counting”) was delivered via MEG-compatible earphones (3M, Saint Paul, MN). The participant closed their eyes and awaited the auditory cue of the scene (e.g. “jungle”) or object (e.g. “bottle”) to imagine, or the number to count in threes from (e.g. “sixty”). The participant then had 3000 ms to imagine or count until a beep indicated the trial was over. The participant then opened their eyes. Seventy-five trials of each condition were presented in a pseudorandom order.

Data were collected using a 275-channel MEG system (CTF, Coquitlam, BC) at a 1200 Hz sample rate, with 3rd order synthetic gradiometry applied. All participants wore three head position coils, placed on the nasion and left/right preauricular points of the head. These coils were energised prior and after each recording block to establish the locations of the sensors relative to these fiducial points.

### 2.2 HCP Data Preprocessing

As the preprocessed data supplied by the HCP consortium is already down sampled to 508 Hz with an anti-aliasing filter applied at ~120 Hz, we opted to process the raw data ourselves to access the higher frequency bands. All preprocessing took place using SPM12 (Litvak et al., 2011). We investigated two different pipelines, where one included an additional denoising step compared to the other. A flowchart of the pipelines is shown in Figure 1 and a detailed description follows below.

**Figure 1.**
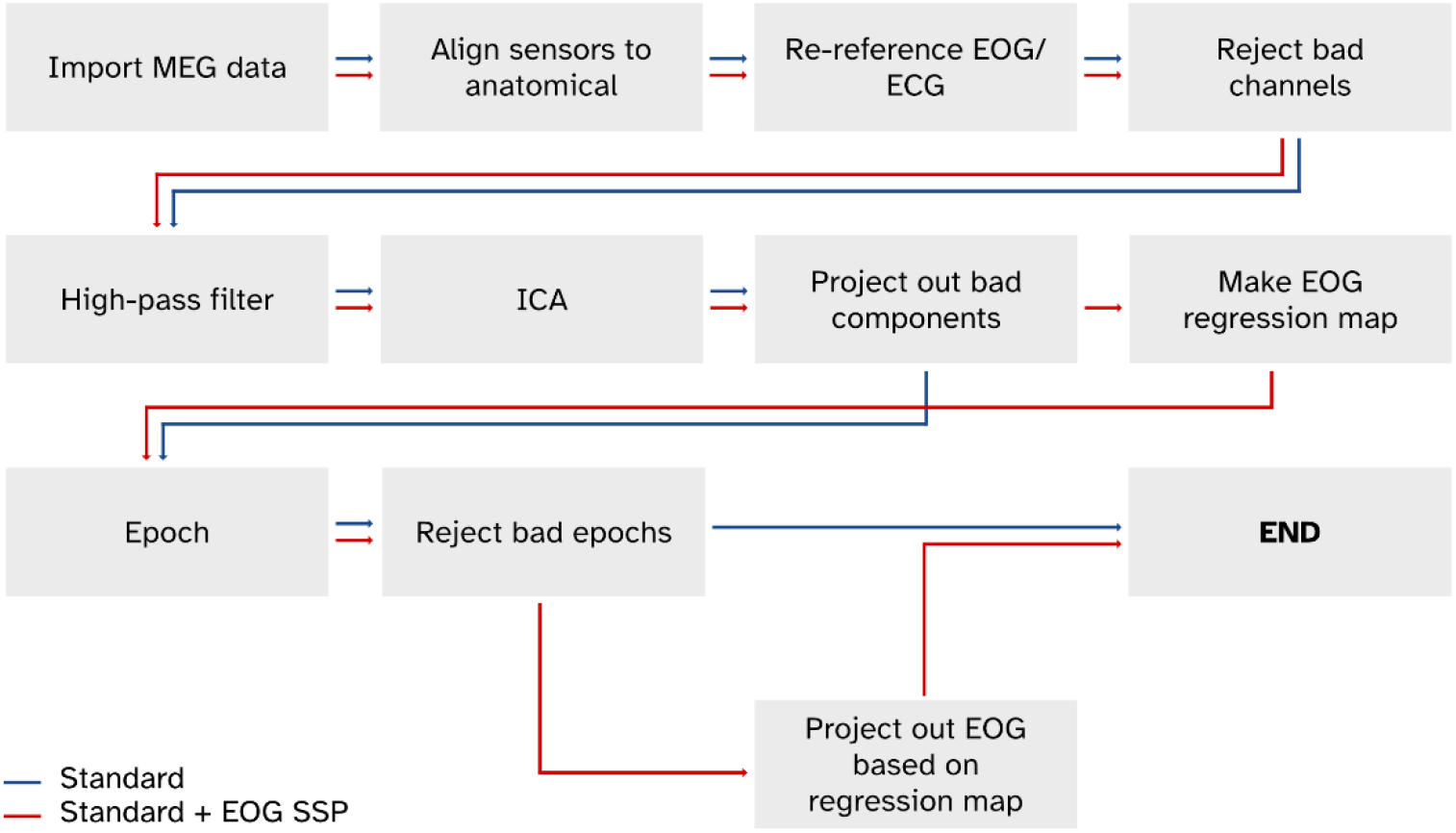
Flowcharts of the two HCP preprocessing pipelines that were deployed. The difference between the standard (blue) pipeline and the red pipeline is the inclusion of an SSP step based on the relationship of the EOG signals and sensor level data for the red pipeline.

#### 2.2.1 Standard Processing (Blue Route)

Data were imported into SPM12 and fiducial locations provided by the HCP consortium were inserted into the dataset for alignment of the sensors with the anatomy. The pairs of H/VEOG and ECG electrodes were subtracted to turn 6 unipolar recordings into 3 bipolar signals. Quality assurance reports from the HCP preprocessed data were used to remove the channels the consortium had already identified as bad, and to identify which periods associated with excessive movement or abnormal EOG/EMG signal to reject later. Next, a 1 Hz high-pass filter was applied to the MEG data.

To denoise the MEG data, we applied temporal independent component analysis (tICA) using FastICA (Hyvarinen, 1999). FastICA was instructed to return 75 components using the symmetric approach. Bad periods in the data (as identified in the HCP QA reports) were masked out prior to component computation. Components which showed strong correlation with the ECG and EOG recordings, as well as components with high kurtosis (which also resembled physiological artefacts) were identified. We used a signal-space-projection (SSP) type approach (Uusitalo and Ilmoniemi, 1997) to remove the offending components, such that for a given instance of time *t,* the MEG sensor level data ***y*** ∈ ℝ^*n_chans_*×1^ can be denoised using

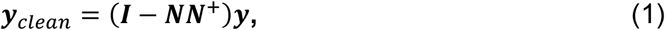

where the ***I*** ∈ ℝ*^n_chans_×n_chans_^* is the identity matrix and the columns of ***N*** ∈ ℝ*^n_chans_×n_comps_^* are the sensor-level topographies of the target components we wish to remove. The superscript + represents the Moore-Penrose pseudoinverse. The dipole forward modelling was updated to account for the gain changes from projecting out the components.

Finally, data were then split into 2 second epochs. Any epoch which intersected a previously-reported bad period in the data were removed.

#### 2.2.2 SSP Projection of EOG Recordings (Red Route)

After ICA, we also investigated the effect of projecting out the high frequency within the EOG recordings from the sensor level data. The horizontal and vertical EOG recordings were bandpass filtered between 100–400 Hz and we fit the signals to the following general linear model:

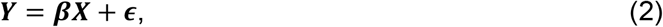

***Y*** ∈ ℝ*^n_chans_×n_samples_^* is continuous sensor-level MEG data, ***X*** ∈ ℝ^2×*n_samples_*^ are the EOG recordings and ***β*** ∈ ℝ^*n_channels_*×2^ are the regression parameters which map the EOG signals to the MEG sensors. These maps were projected out using the SSP framework described in Equation 1 once the data had been epoched and bad trials rejected. Again, compensation matrices were updated to reflect the subspace.

### 2.3 Scene Imagination Data Preprocessing

Preprocessing of the scene imagination data followed a similar but not identical pipeline. First, a visual inspection of the data to identify SQUID resets and jumps, with trials containing these omitted from further analyses. Subject-specific anatomical images were not present with this dataset, so a template anatomical based on the MNI-152 standard was registered, where a 7 degrees of freedom affine transformation was used to match the anatomical fiducials the locations of the HPI coils on the subject. Data were then filtered using a high-pass filter with a 1 Hz cut-off band and tICA using the FastICA algorithm was performed, again returning 75 components. Components representing unwanted components (blinks, saccades and ECG waveforms) were projected out using the SSP framework outlined in Equation 1. No projection of EOG channels were performed due to a lack of EOG recordings in the dataset.

### 2.4 Source Analysis

All source analysis was performed with the DAiSS toolbox within SPM12. After the anatomical images were registered with the MEG data, a head model representing the boundary mesh between the cerebrospinal fluid (CSF) and the inner skull was generated. Subsequently a 5 mm volumetric grid of sources within this mesh was generated. Three orthogonal current dipoles at each location were modelled using Nolte’s single shell method (Nolte, 2003) and reduced in rank to 2 principal orientations. Epoched sensor-level data were discrete-cosine-transformed (DCT) to filter into a band of choice and covariance matrices for each trial were generated, then summed across epochs. Regularisation of the covariance was performed by truncating the total number of principal components to 200 for the HCP data and 250 for the scene imagination data, which were sufficiently before the cliff-face in the eigenvalues due to removal of interference data (Westner et al., 2022). Source reconstructions were performed with a Linearly Constrained Minimum Variance (LCMV) beamformer (Brookes et al., 2008; Van Veen et al., 1997), where the three components where linearly combined for maximal reconstructed variance (Sekihara et al., 2004).

### 2.5 Power Analysis

For the HCP dataset, depth-corrected and noise-subtracted source power for the *i*^th^ location was calculated using

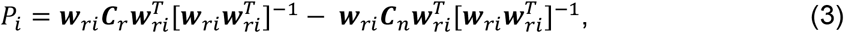

where ***w**_ri_* ∈ ℝ^1×*n_s_*^ represents the resting state source reconstruction weights from the *n_s_* sensors to source *i, **C**_r_* ∈ ℝ*^n_s_×n_s_^* is the DCT-filtered sensor covariance matrix from the resting state data and ***C**_n_* ∈ ℝ*^n_s_×n_s_^* is the covariance from the empty room recording. No additional smoothing of the images was applied. For each resultant image, the top 1800 voxels (where Eqn. 3 was positive, i.e. projected brain power estimates were greater than projected empty room estimates) were identified and binarised to make a peak ROI. The choice to threshold the top 1800 voxels (i.e. 0.75% of the brain volume) is arbitrary, but will determine the probability of overlap across subjects. Based on this threshold we ran multiple simulations at different levels of smoothing to determine how likely it would be for the same supra-threshold voxel to overlap by chance across multiple subjects. For 89 images, we found overlap in 8 subjects or more in a given voxel would have a probability of p < 0.05 that it could be explained by images generated of random noise (see supplementary information for details of the calculation).

For the scene imagination data, there were no associated noise recordings collected with the subject data, so power images could not be noise-subtracted. Here the source power for the *i*^th^ was therefore calculated as

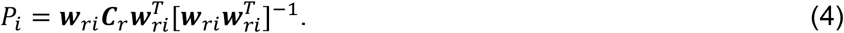

Furthermore, with just 22 subjects, our threshold for the number of subjects have peak voxels significantly overlap is 5.

### 2.6 Functional Connectivity Analysis

Functional connectivity can be measured in a multitude of ways which look at phase-based relationships (Lachaux et al., 1999; Stam et al., 2007; Vinck et al., 2011), amplitude-based (Hipp et al., 2012; O’Neill et al., 2015) or even a combination of the two (Canolty et al., 2006). Here we opted for amplitude envelope coupling between two regions due to its robustness, particularly within the HCP dataset (Colclough et al., 2016). Virtual electrodes were derived from the centroids of the AAL3 parcellation (with the cerebellar regions of interest omitted), leaving 90 areas of interest. Virtual electrodes were then band pass-filtered into a band of choice. Leakage correction was performed using symmetric orthogonalization (Colclough et al., 2015), and the amplitude envelopes of the signals were derived via Hilbert transform. The resultant envelopes were then correlated to generate a connectome per subject.

### 2.7 Coherence Analysis

Coherence measures between the re-referenced EOG recordings and the source reconstructed MEG data were calculated using the Dynamic Imaging of Coherent Sources method (DICS; Gross et al., 2001) using the existing LCMV beamformer weights. Note that this analysis was only performed on the HCP data.

### 2.8 Data and Code Availability

MEG data from the Human Connectome Project is freely available as part of the HCP1200 data release from https://db.humanconnectome.org. An account needs to be registered prior to access. The scene imagination dataset is available upon reasonable request – please contact EAM. An archive of the processing scripts used for this study are available at https://github.com/georgeoneill/study-2023-hfa.

## 3. Results

The results presented in sections 3.1–3.4 relate to the HCP resting dataset, while the results for the scene imagery data in section 3.5.

### 3.1 Localisation of Source Power

Figure 2A is a set of glass brain plots which contain the conjunction of peak power across all 89 subjects in the 100–200 Hz, 200-300 Hz and 300-400 Hz bands, with our standard preprocessing applied prior to source localisation. We only show the voxels where the overlap across subjects was 8 or larger (p<0.05). In all bands we see that power is primarily distributed in both anterior temporal lobes, with a preference for the right hemisphere.

**Figure 2:**
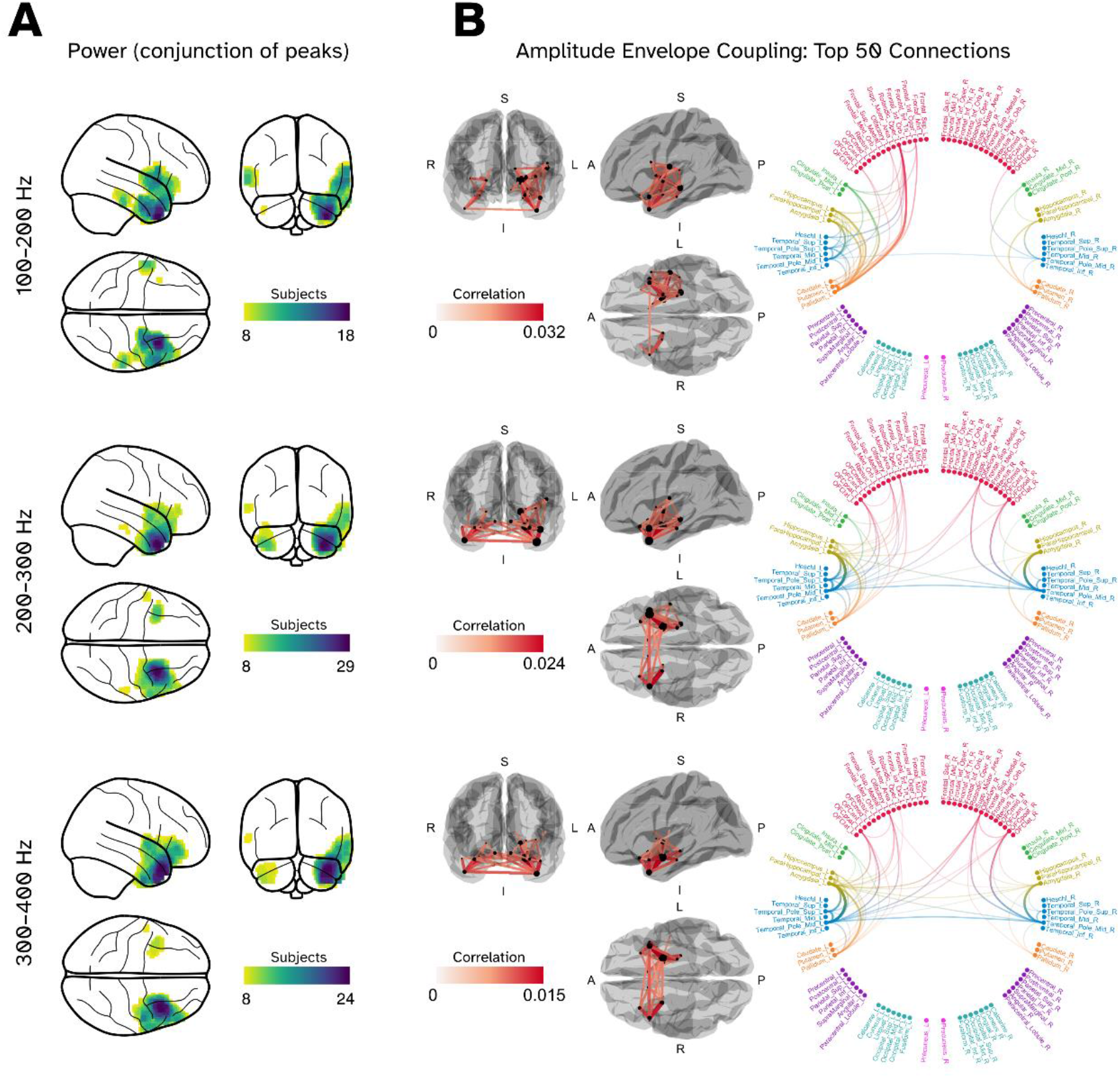
A robust network found in resting state MEG data (n=89) in bands above 100 Hz. (A) Glass brain plots mapping the conjunction of peak power across the subjects in the 100–200 Hz (top panel), 200–300 Hz (middle panel) and 300–400 Hz (bottom panel) bands. Only significant (p<0.05) overlap of 8 subjects or more are shown. (B) Subject averaged amplitude envelope coupling connectomes displayed in a 3D plot and a chord plot respectively.

In the 100-200 Hz band (Fig. 2A, top panel), peak conjunction occurred in the *right temporal pole* (18 subjects; MNI location: [28 8 −40] mm) with separate clusters centred over the *right insula* (16 subjects; [56 10 0] mm), *right inferior temporal lobe* (11 subjects; [52 −28 −32] mm), *left insula* (10 subjects; [−62 −4 0] mm) *left temporal pole* (8 subjects; [−40 5 −42] mm). Investigating the 200–300 Hz band (Fig. 2A, middle panel) we see that largest overlap occurs in the right temporal pole, with 29 subjects overlapping ([32 12 −42] mm). We also see an increase in subjects overlapping in the left temporal pole (14 subjects), and a cluster in the right insula (peaking at 13 subjects). Also apparent is a similar distribution of peaks in the 300-400 Hz band (Fig. 2A, bottom panel).

### 3.2 Amplitude Envelope Coupling

Subject-averaged functional connectomes from the data are shown in Figure 2B which, for clarity, are only showing the top 50 connections (by average correlation). We observed that the areas which showed the largest areas of power overlap in Figure 2A are largely employed in the functional networks observed. In the 100-200 Hz band our strongest connections exist between the left temporal pole and the amygdala nodes, as well as strong connections between areas such as the left hippocampus, left insula and left pallidum. We see a similar, but weaker, set of connections mirrored in the right hemisphere. We now also see a single interhemispheric connection linking the temporal poles make it into the top 50 connections. 200-300 Hz, the strongest connections, are again between the temporal pole and amygdala, but this time the connections are similar in strength in both hemispheres. The number of interhemispheric connections in the top 50 here also increase to 6; the temporal poles are connected to each other, as well as the amygdalae, while we also see the right olfactory cortex connected to nodes in the other hemisphere. Our connectivity patterns in the 300-400 Hz band is similar in topography, but with increasing interhemispheric connectivity (10/50 connections cross the midline).

### 3.3 Relationship Between EOG Signals and the Brain

We investigated further the relationship between the EOG signals and sources in the brain in the HCP dataset. Figure 3A shows the subject-averaged coherence between the HEOG (top panel) and VEOG (bottom panel) within the 100–200 Hz band prior to the EOG signals being projected out. We observed that the HEOG signal is most coherent in the anterior temporal lobes and temporal poles, with stronger coherence in the right hemisphere compared to the left. The VEOG coherence, whilst again stronger in the right hemisphere, is localised to the inferior frontal lobe. We also note that the coherence is approximately a third of the intensity compared to the HEOG. Figure 3B shows the subject-average regression maps between the MEG sensors and the EOG signals prior to SSP being applied between this time between 100–400 Hz. The HEOG map has two poles across the anterior temporal sensors, but there is a large variability across subjects, and does not strongly resemble the sensor-level map of saccadic spikes as shown in (Carl et al., 2012), whereas the VEOG map has the strongest regression artefacts in channels which are associated with the blink artefact (Hari and Puce, 2017). However, it is interesting to note that the topography appears to resemble anterior, bilateral dipoles with opposite orientations. Finally, Figure 3C depicts the relationship between the variance in the EOG signals and the peak beamformer power in the anterior temporal lobes. Here the ROIs selected were derived from the anterior temporal lobe parcels from the Schafer 200 atlas (Lawrence et al., 2021; Schaefer et al., 2018). We see that prior to SSP, there was a slight but significant linear relationship between the power estimate at the anterior temporal lobe and the HEOG variance (R^2^=0.077; F=7.097; p=0.009). Applying SSP removed the significant relationship (R^2^=0.038; F=3.181; p=0.079). For the VEOG signals there was no significant link to the beamformer power prior to or after SSP (p=0.301 v p=0.401). The detailed results for the 200–300 Hz and 300–400 Hz bands are in the supplementary material, but in short, applying SSP breaks the significant relationship between the anterior temporal lobes and the HEOG variance in the 200–300 Hz band, but not in the 300–400 Hz band.

**Figure 3:**
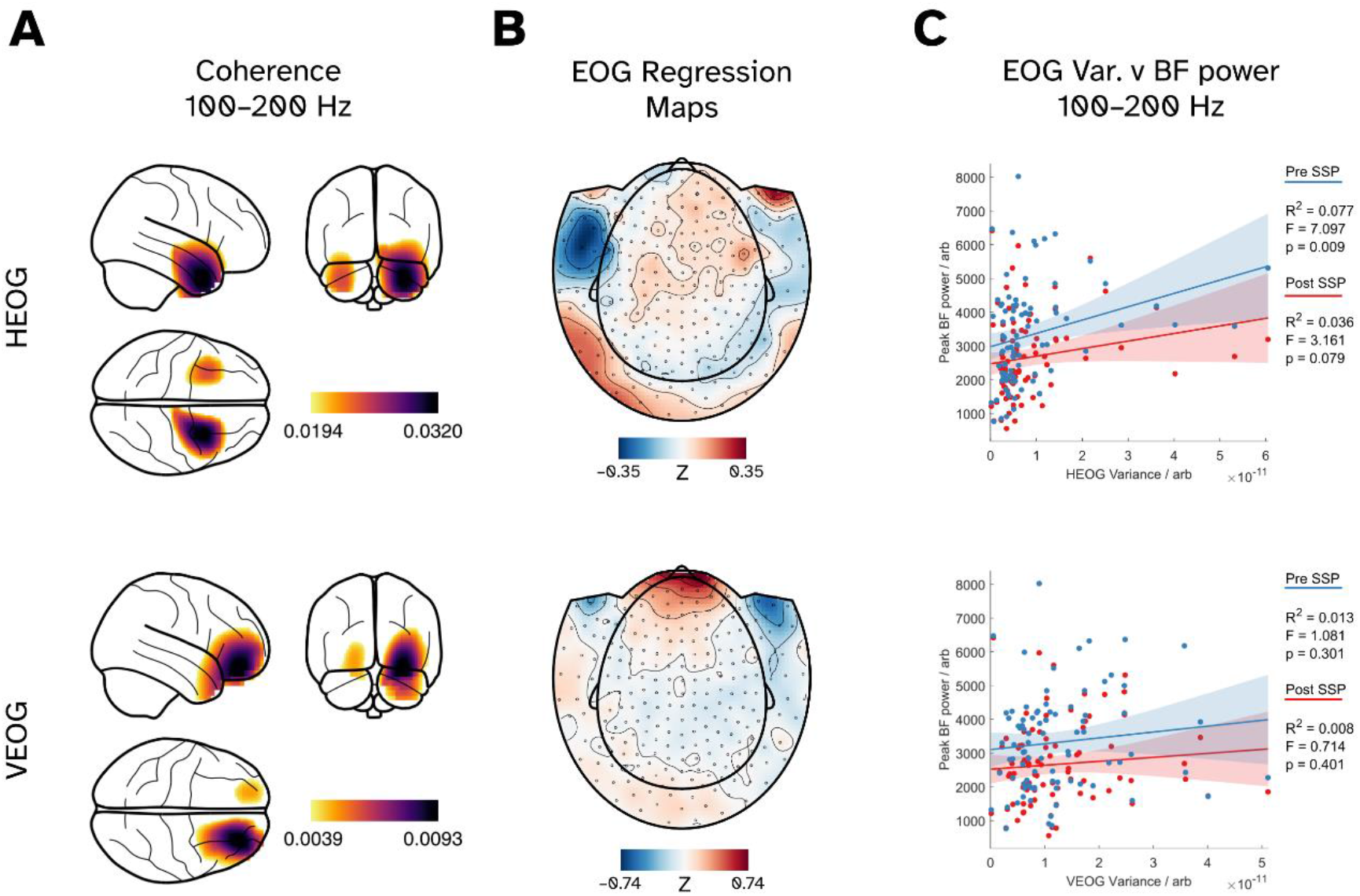
Investigating the relationship between the EOG and MEG signals in source and sensor space. The top row relates to the horizontal EOG (HEOG) signals whilst the bottom row relates to the vertical (VEOG). (A) Glass brain plots of the coherence between EOG and reconstructed sources in the brain prior to any SSP of the EOG signals being applied. (B) The subject-averaged regression maps between the MEG sensors and EOG signals. The individual maps are used for SSP later. (C) Scatter plots showing the relationships between the variance of the EOG signal and the maximal beamformer power reconstructed in the anterior temporal lobe before (blue) and after (red) projection of the EOG signals.

### 3.4 The Effect of Projecting Out the EOG Signals from the Data

Given the robust coherence between the sources in the temporal lobes and the HEOG recordings, we further investigated how additional denoising by projecting out the EOG data affected our initial results.

Figure 4 shows our power conjunction results, similar to Figure 2, but this time using the data after an EOG SSP has been applied. Again, we only show voxels which met our significance threshold (p<0.05) of 8 subjects or higher overlapping. Below each conjunction plot is a map showing the difference in the number of subjects which overlap when compared to our standard processing pipeline (which does not include removing the EOG signals). Applying an EOG SSP denoising in the 100–200 Hz band attenuated the peaks in the right temporal pole (overlap dropped from 18 subjects to only 4, below our significance threshold), the left temporal pole drops from 8 subjects to 6 in comparison, and other noted peaks remain above our critical thresholds. In the higher bands, we also see our region of interest in the right temporal pole is heavily affected, with a reduction of 29 to 10 subjects overlapping in those areas between 200–300 Hz and from 24 to 9 between 300-400 Hz. Focusing on 200–300 Hz, a second peak in the RTP is revealed with 17 subjects; the peak is marginally posterior compared the previous peak ([40 2 −44] mm versus [32 12 −42] mm). We can see clearer clusters of overlap in the right superior temporal lobe (17 subjects; [60 2 −4] mm) and an equivalent in the contralateral hemisphere (11 subjects; [−60 −8 −4] mm), with the maximal overlap of subjects in the left temporal pole also being 10. An interesting feature in the 300–400 Hz band is the large increase in overlap of subjects in the right inferior frontal/pars triangularis area (3 to 17 subjects; [52 38 0] mm).

**Figure 4:**
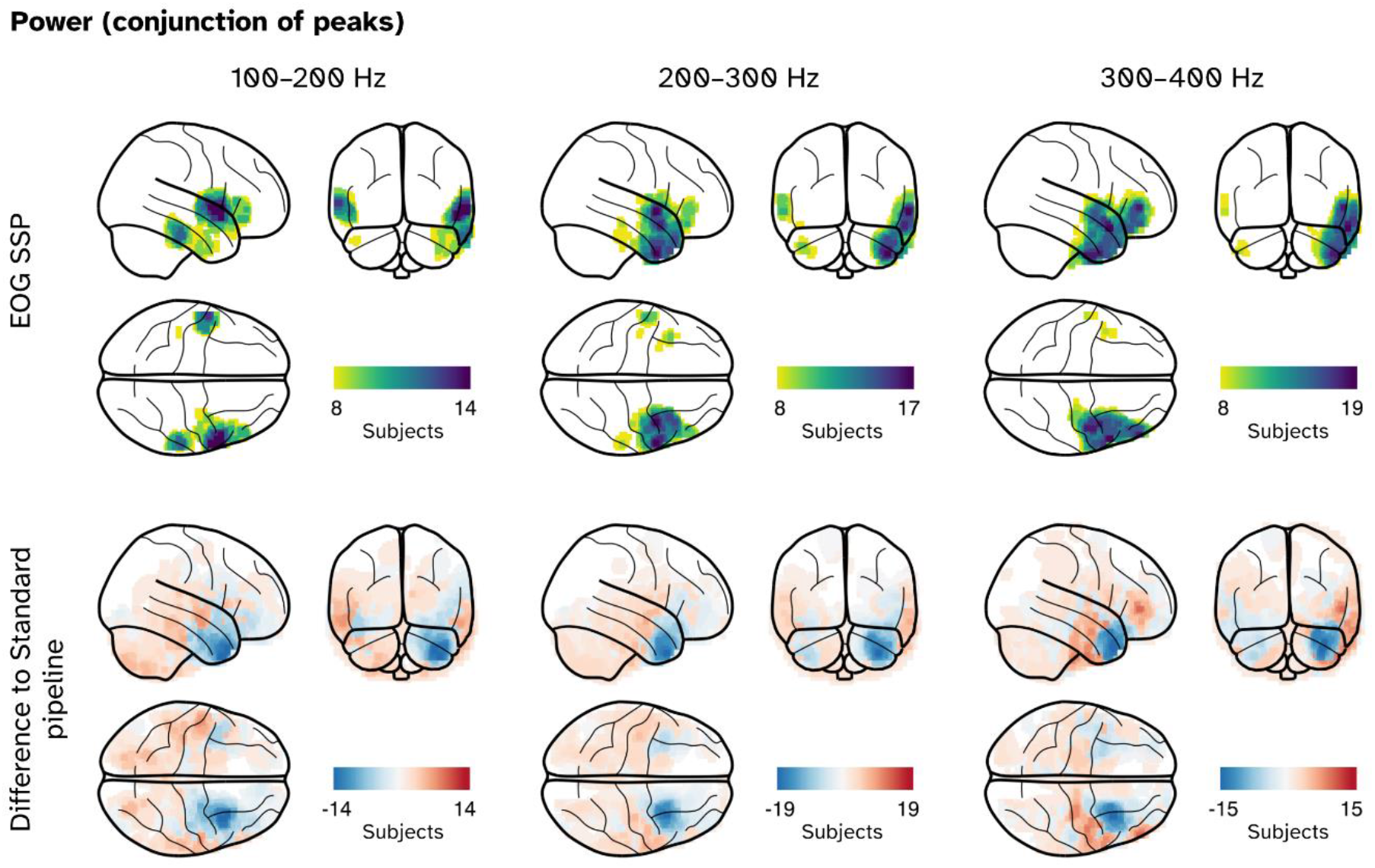
Conjunction maps of peak power, in this case for data where an additional denoising step was included, where EOG signals were projected out of the MEG data. Top row: The resultant conjunction maps, where the number of subjects overlapping in a given voxel reached or exceeded our critical (p < 0.05) threshold of 8. Bottom row: difference in the number of overlapping subjects between the datasets without EOG signals removed (Figure 2) and with.

The effects of applying the EOG SSP to the connectivity results can be seen in Figures 5 and 6. Focusing on 100–200 Hz data in Figure 5, we see the top 50 connections look similar (Fig. 5A), with 44 connections being shared between the pre-SSP and post-SSP connectomes. Whilst the topology of the networks between preprocessing methods were similar, there was a moderate reduction in connectivity after SSP has been applied (peak connectivity reduced from 0.032 to 0.028). Paired-T tests between each connection showed the reduction in connectivity as significant for 254/4005 node pairs (p < 0.05, FDR corrected; q=0.05). A map of these significant changes is found in Figure 5B, but contains primarily fronto-temporal areas. Within the union of each set of top 50 connections, 27 showed significant differences in connectivity. The results are depicted in Figure 5C, which shows the connections in the left hemisphere (as well as the sole interhemispheric connection in the list) primarily affected.

**Figure 5:**
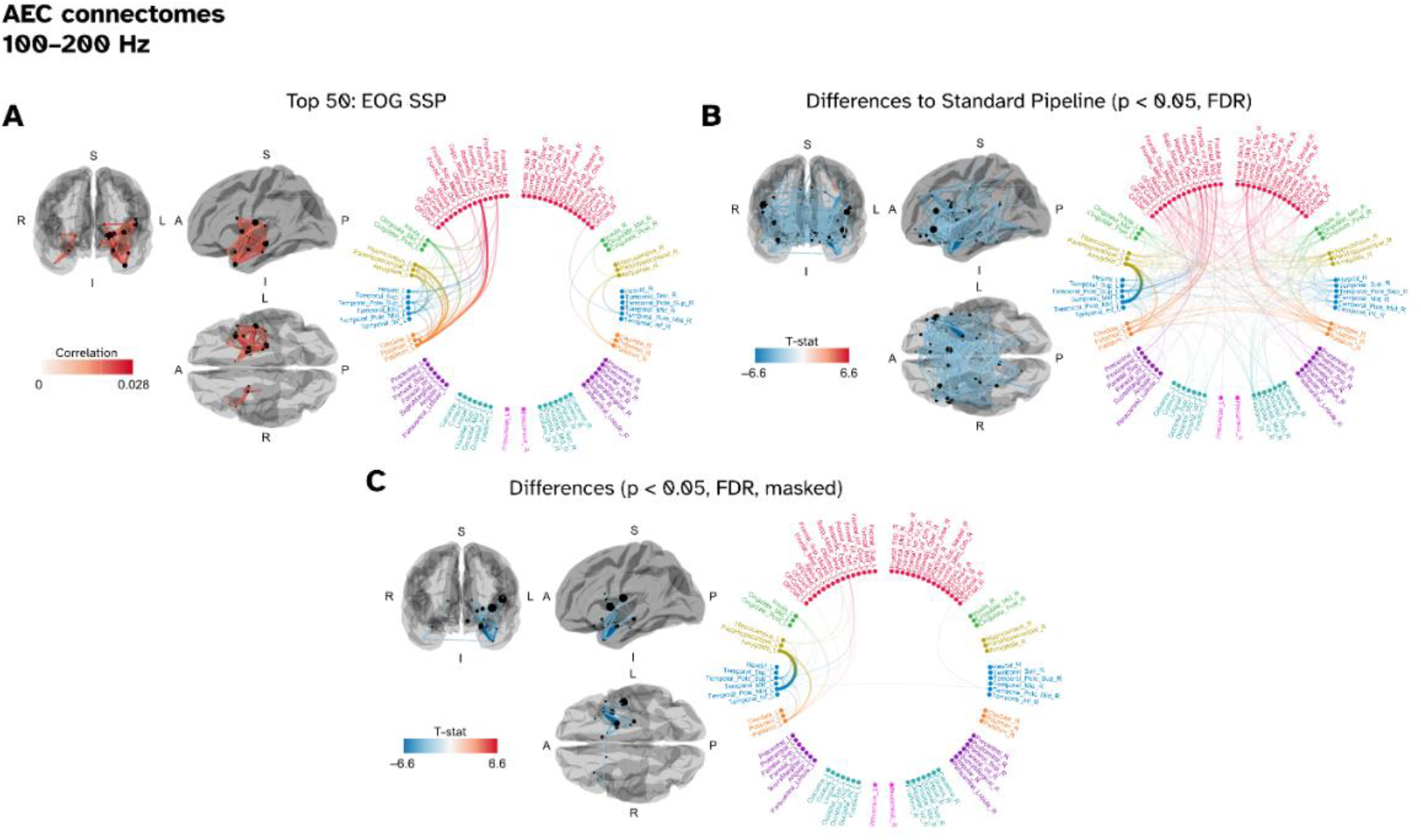
The effect of projecting out the EOG signal on the functional connectome in the 100–200 Hz band. (A) The top 50 connections of the subject-averaged connectome, as defined using the amplitude envelope coupling metric, showing a similar network of connections to the pre-EOG-projected results in Figure 2. (B) Results of paired-T tests between the pre- and post-projected connectomes. Plotted connections show the significant (p < 0.05, FDR corrected) differences, revealing a widespread reduction in connection strength in fronto-temporal areas. (C) Significant connections but this time masked to within the top-50 connections.

**Figure 6:**
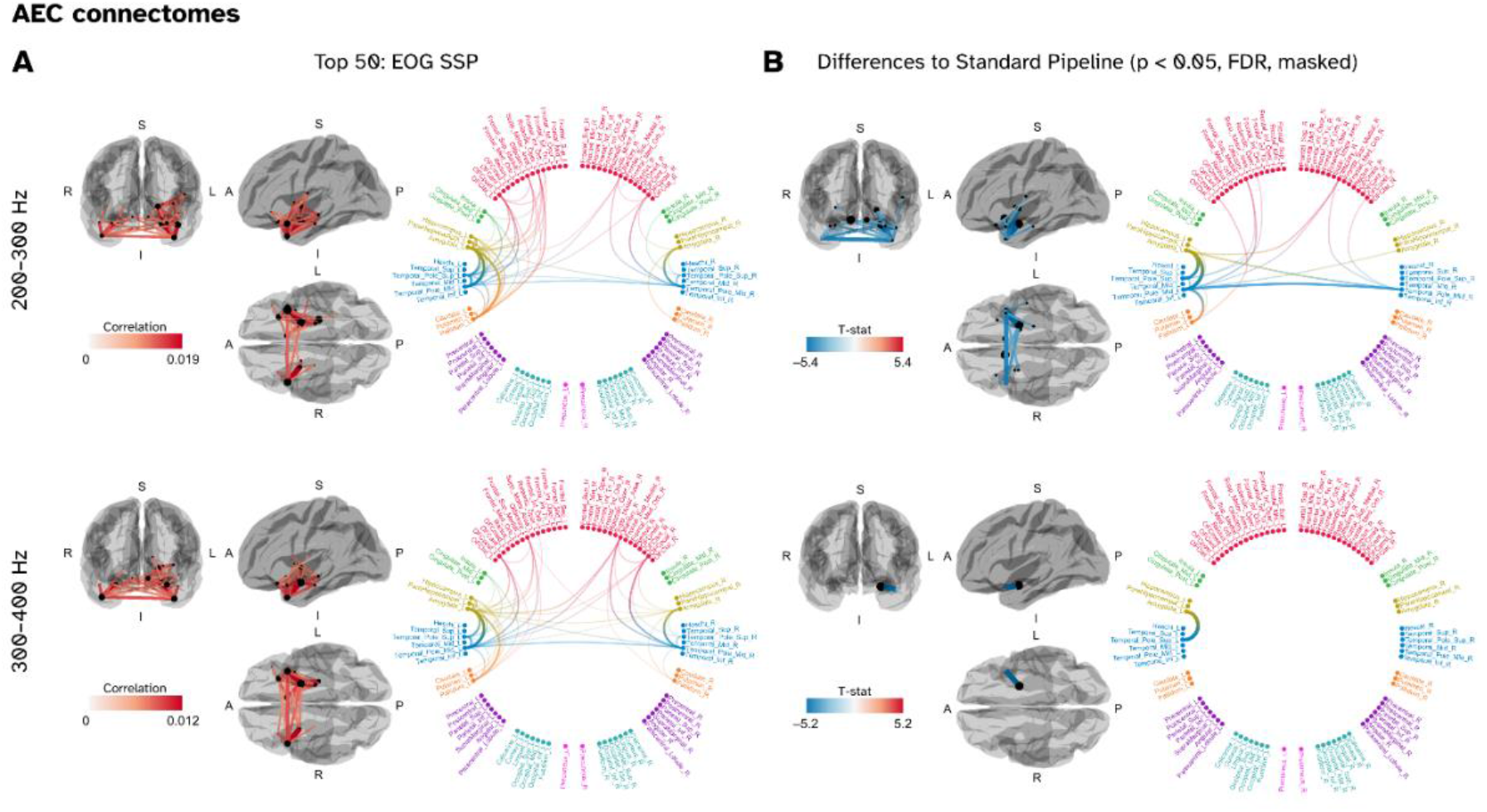
The effect of projecting out the EOG signal on the functional connectome in the 200–300 Hz and 300–400 Hz bands. (A) The top 50 connections of the subject averaged connectome. (B) The significant differences significant (p < 0.05, FDR corrected) compared to the pre-projected data, masked within the top 50 connections for clarity.

We see a similar effect in the 200–300 Hz and 300–400 Hz connectomes when the EOG data are projected out. This is depicted in Figure 6. Again, the top 50 connections in each band were highly similar to the non-EOG-SSP counterparts (44/50 shared for 200–300 Hz and 44/50 for 300–400 Hz). Peak connectivity was again reduced, with paired-T tests showing in the 200–300 Hz band many of the connections were significantly reduced in magnitude (Fig. 6B), with a particular note that most of the interhemispheric connections were included. In the 300–400 Hz band only a single connection in the whole brain was deemed as significantly reduced after false discovery correction – the link between the left amygdala and the left temporal pole.

### 3.5 Replication of network in a task-based dataset

We wanted to investigate whether the features we saw in the HCP data would be observable in a different dataset, so applied our analyses to the novel scene imagination task dataset (Barry et al., 2019). The results of the power conjunction analysis are depicted in Figure 7A, where based on our especial testing an overlap of 5 subjects or more in a given voxel was considered significant (p < 0.05). In the 100–200 Hz band we have only two significant clusters, in bilateral superior temporal gyri (MNI locations [60 −10 0] mm and [−54 −4 −10] mm). At 200–300 Hz we also observe two significant clusters in the cortex, one in the left superior temporal gyrus ([−62 −8 −10] mm) and the left temporal pole ([−44 4 −40] mm). Note that due to the CTF MEG system applying a hardware anti-aliasing low-pass filter at 300 Hz, we do not have the results for the 300–400 Hz band. The functional network analysis is depicted in Figure 7B, there the top 50 strongest connections (defined by amplitude envelope coupling) are shown. Similar to the networks seen in the HCP dataset we see our strongest (within hemisphere) connections in the 100–200 Hz band between the hippocampal, amygdalae and temporal pole areas. We also observe similar networks to the HCP in the 200–300 Hz band, with increasing number of interhemispheric connections also reported.

**Figure 7:**
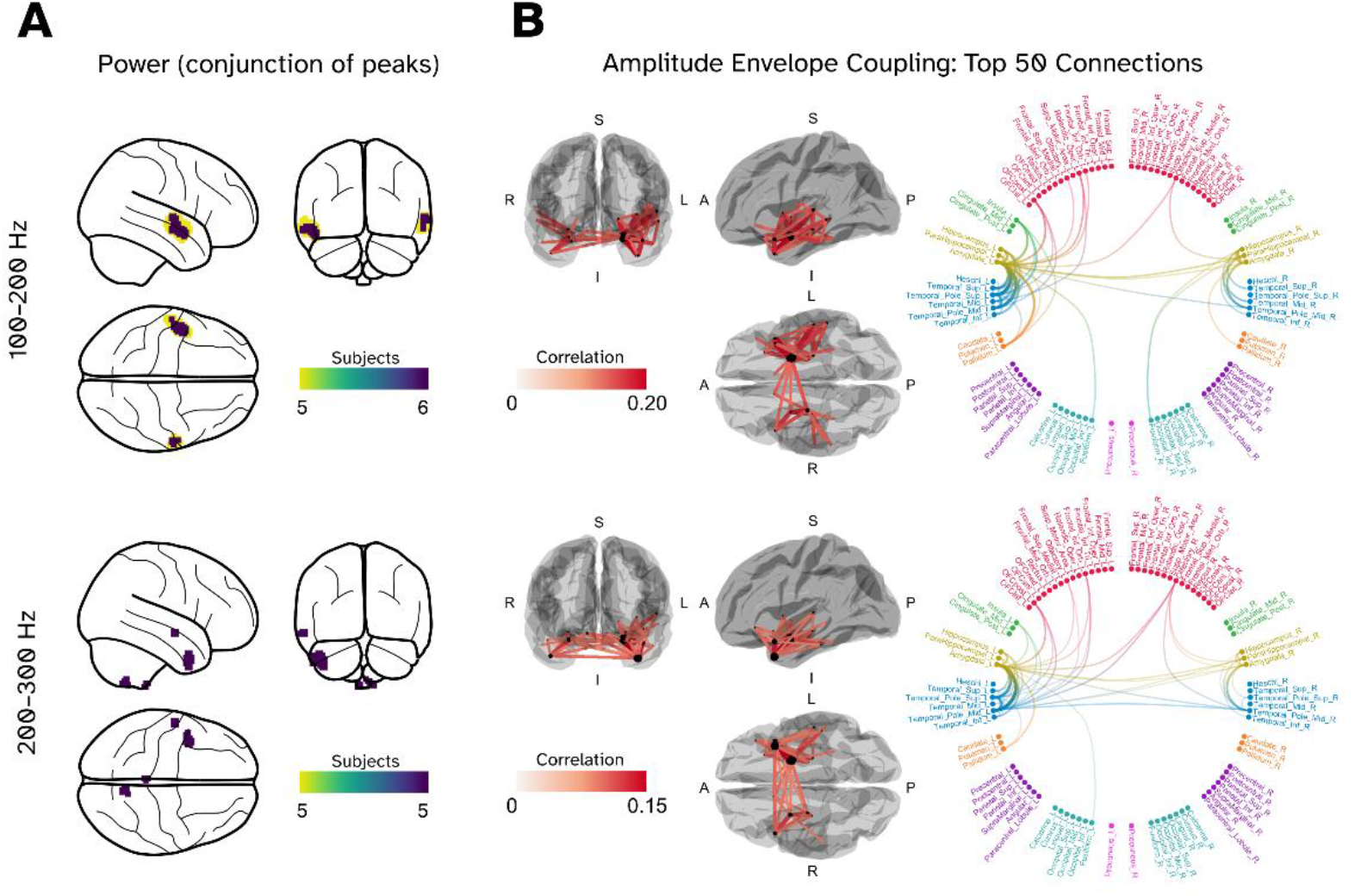
Replication of the high frequency functional network in a second, task-based, dataset with 22 subjects (Barry et al., 2019). (A) Power conjunction maps within the 100–200 Hz (top panel) and 200–300 Hz (bottom panel) bands. The displayed voxels where the number of subjects overlapping in a given voxel reached or exceeded our critical (p < 0.05) threshold of 5. (B) Left column: glass brain plots of the top 50 connections in the subject-averaged connectomes for the 100–200 Hz (top panel) and 200–300 Hz (bottom panel) bands. Right column: Associated chord plots showing which nodes were employed in these 50 connections, thicker lines indicate stronger coupling.

## 4. Discussion

We have shown that conventional MEG analysis of eyes-open resting state data gives rise to an apparent high-frequency anterior temporal network which is reproducible across 100 Hz bands up until 400 Hz. These networks appear to be reproducible – not only did we observe this in the Human Connectome Project resting state data, but similar topographies were also found in a 22-subject, task-based dataset collected in a previous study (Barry et al., 2019; O’Neill et al., 2021) that employed a different MEG system in a different country. Using the complimentary empty-room recordings of the HCP dataset we cannot simply replicate the network topographies without recording from the brain of a person within the MEG scanner (supplementary material). It appears to be a genuine feature in the MEG data which is (at minimum) of biological origin and not the product of instrumental or environmental noise. We posit that these features will be common to all recorded human MEG data.

However, we are unable to definitively answer whether these signals are of neuronal or myogenic origin. Using our peak localisation method to identify where beamformer power was being regularly localised, we saw that power was clustering increasingly anterior within the temporal lobe the higher we progressed through the bands. Furthermore, the bilateral functional connections increasing in representation in a similar fashion suggests that we might have been mis-localising bilateral magneto-ocular artefacts within the temporal lobe. Coherence maps between the EOG data and these resting state images show the highest coherence in key anterior temporal and anterior frontal regions (Figure 3), lending credence to this idea. We know from the previous literature that even highly targeted invasive recordings in the anterior temporal lobe can still detect the oculargram being conducted into the brain tissue (Jerbi et al., 2009).

After projecting out EOG measured activity from the MEG data, not only did we see the peaks overlap less in the temporal poles (Figure 4) but we could also break the significant linear relationship between the horizontal EOG variance and beamformer power localised to the anterior temporal lobes (Figure 3). However, looking at connectivity measures, there remains an anterior temporal network, operating at over 100Hz. Whilst the raw connectivity did show significant reduction in global coupling, ultimately the topology of the network has not been altered (Figures 5 and 6).

A parsimonious explanation is that these data derive entirely from eye muscle, some components of which were not picked up on the EOG recordings. The signals are not only high-frequency, but also broadband – consistent with previous reports of muscle activity (Jerbi et al., 2009; Muthukumaraswamy, 2013) and a recent study which indicates a strong relationship between EEG data and EOG recordings high frequencies (Quinn et al., 2022). The location, at the anterior temporal pole, is also consistent with the location of the lateral rectus muscle, previously identified as a source of high-frequency electrical activity (Carl et al., 2012; Jerbi et al., 2009). However, the magnetic field profile of this activity (Figure 3B) does not match up with the saccadic spike artefact as measured with MEG and EEG by Carl et al. (2012).

A more complex yet also potentially more exciting explanation is that in addition to eye-muscle activity, a high-frequency cortical network is in operation. This operation is perhaps related to the default mode network (Higgins et al., 2021) and/or continuous or ongoing processing associated with memory replay (Liu et al., 2019). Still more complex is that the EOG signals we had access to derived from the brain rather than the muscle or eyeball. One confusing factor is that the lateral rectus muscles operate in opposition – so one would anticipate anti-correlated high frequency envelope activity between left and right temporal lobe sources. However, we see no evidence for this in our connectivity results, where the envelope coupling in the temporal poles was positively correlated, but we may possibly see hints of this in the VEOG sensor map (Figure 3B), which appears to show the topography of two dipoles in opposite orientations. It is also noteworthy that similar confounds/questions exist for the vertical EOG channels and inferior frontal cortex, where we again see coherence between the two measures. However, we also note the coherence was considerably lower than for the horizontal EOG, so the effect may be less pernicious.

Independent component analysis (ICA) as a means to remove unwanted features in M/EEG data has been an established strategy and is often recommended as an essential step in data preoprocessing (Gross et al., 2013). Indeed, this study implemented ICA as an attempt to separate ocular signals from the neural data, but it appears this alone is not entirely effective in removing these signals. From our experiences, we can identify three potential points of failure. First, is the choice of ICA implementation itself. In particular we used FastICA (Hyvarinen, 1999) which, whilst a powerful tool for removing vertical EOG artefacts (such as blinks), is suboptimal at removing saccades typically associated with horizontal eye motion (Mäkelä et al., 2022). The authors of the aforementioned study found the ability to remove saccadic artefacts was improved when using it in conjunction with a second algorithm or replaced entirely. Their recommendation is to use the AMICA method (Palmer et al., 2008), which does not assume sources are stationary. The second issue is the robustness at identifying which components are artefactual, which will only be as good as the persons reviewing the components. Heuristics such as the correlation of components to the EOG/ECG signals can help with this, but are not always present. Helpfully, methods which use machine learning methods to identify the origin of a component are starting to become available for MEG data (Treacher et al., 2021). Finally, a third potential issue is the assumption the spatial topographies of these artefacts are similar across all frequencies. We used broadband data (1–400 Hz) to generate the component topographies we wanted to project out of the data, which assumes (for example) a blink artefact which dominates the spectrum up to ~5 Hz (Keren et al., 2010), will share the same spatial profile as signals over 100 Hz. This may not be an effective approach and what would be more helpful in the future is to perform ICA on data filtered into the high frequency bands of choice.

To conclude, we remain excited that these high-frequency signals could be of neuronal origin but simply caution that we cannot rule out that they are generated by eye muscles. Our sole recommendation for future studies would be to use some independent metric of this activity (EOG/optical eye tracking or both) when focusing on higher-frequency regions of the spectrum.

## 5. Acknowledgements

Funding for this project was provided by EPSRC (EP/T001046/1) from the Quantum Technology hub in sensing and timing (sub-award QTPRF02). SM is funded by EPSRC-funded UCL Centre for Doctoral Training in Medical Imaging (EP/L016478/1) and the Department of Health’s NIHR-funded Biomedical Research Centre at University College London Hospitals. NA, RAS and EAM are supported by a Wellcome Principal Research Fellowship awarded to EAM. (210567/Z/18/Z). TMT is funded by a fellowship from Epilepsy Research UK and Young Epilepsy (FY2101). The Wellcome Centre for Human Neuroimaging is supported by core funding from Wellcome (203147/Z/16/Z).

We thank Daniel Barry for his work on the original scene imagination MEG study.

