## Supplementary Material for "Is high-frequency activity evidence of an anterior temporal lobe network or micro-saccades?"

George C. O'Neill[a*], Stephanie Mellor[a], Robert A. Seymour[a], Nicholas Alexander[a], Tim M. Tierney[a], Ryan C. Timms[a], Eleanor A. Maguire[a], Gareth R. Barnes[a]

[a]*Wellcome Centre for Human Neuroimaging, Department of Imaging Neuroscience, UCL Queen Square Institute of Neurology, University College London, London, UK*

---

***1) Deriving the empirical lower bound for subject overlap.***

In the main manuscript we derived an empirical result for 89 subjects whereby an overlap of 8 binarised images containing 1800 voxels exceeded the null threshold of $p < 0.05$. To reach that conclusion we performed the following:

Our null hypothesis was that thresholding and binarising random noise could equally generate the same number of voxels to overlap for a given number of subjects. Hypothetically for data where there is no spatial autocorrelation between voxels, it is entirely plausible that a binomial distribution could tell us where the statistical limit would exist. However, we necessarily assume our data is smooth, so this likely gives us too low an estimate as to where the real threshold should lie. We could use random field theory (RFT; Worsley et al., 1996) to find this limit, but it is known that images generated from beamformers have differing levels of smoothness within a single image (Barnes and Hillebrand, 2003), and so assumptions about RFT are violated. Instead, we opted to sample a threshold from thousands of random fields with varying smoothness to find our limit.

For a given target smoothness, 1000 volumetric images of dimensions 273 × 327 × 273 (3 times the size of a standard MNI-152 image with 2mm resolution) were generated. Then if required, a spherical gaussian smoothing kernel was applied to the noise. We opted for such a large image such that edge effects of the smoothing would not affect the centre of the image volume. Next, a brain mask from the MNI-152 brain sitting in the centre of the volume was applied to all images and then thresholded so only the top 1800 voxels survived. The surviving voxel locations were stored for each of the 1000 images.

The next step was to generate the null distribution from the bank of noise images. Here, for a given iteration of the null, 89/1000 images were randomly selected, and the list of voxels concatenated. The number of times the mode voxel appeared in the list was counted to build up a null distribution. 500 iterations of random selection for a given number of subjects and smoothness were used to build the null distribution, and the $95^{th}$ percentile was used to set the limit.

The results of this can be found in Figure S1. Figure S1A shows the results for assuming all voxels are uncorrelated (no smoothing). In blue is the histogram of how many voxels in the brain mask overlap when 1000 images are generated, the median response is 7 and the maximum is 23. Sampling 89 images from the bank of 1000 images and finding the max overlap multiple times generates the distribution in shown in orange. Here, for this example we see that the $95^{th}$ percentile happens at 8 images overlapping. We also note that this exceeds what a binomial distribution (89 trials with a probability of 0.0075 – the number of binarised voxels in our brain mask) would set the $95^{th}$ percentile, which would be 3. Figure 1B shows how the threshold for 89 subjects is affected by smoothing. We used smoothing kernels with 1, 2, 3, 4, 5, 8, 10, 12, 15, 20, 25, 30, 35 and 40mm full width half maxima. Here we see that as the smoothing becomes more extreme the number of overlaps starts to drop, in this case from 8 to 7. Finally, we investigated how the number of subjects for a given smoothness affects the threshold. The results with a 5 mm kernel and a number of subjects logarithmically spaced between 1-1000 is shown in Figure 1C. Our initial hypothesis was that the threshold should approximate the square root of the number of subjects, and that is shown with a black dashed line. It appears to hold for low (<100) subjects but deviates above that.

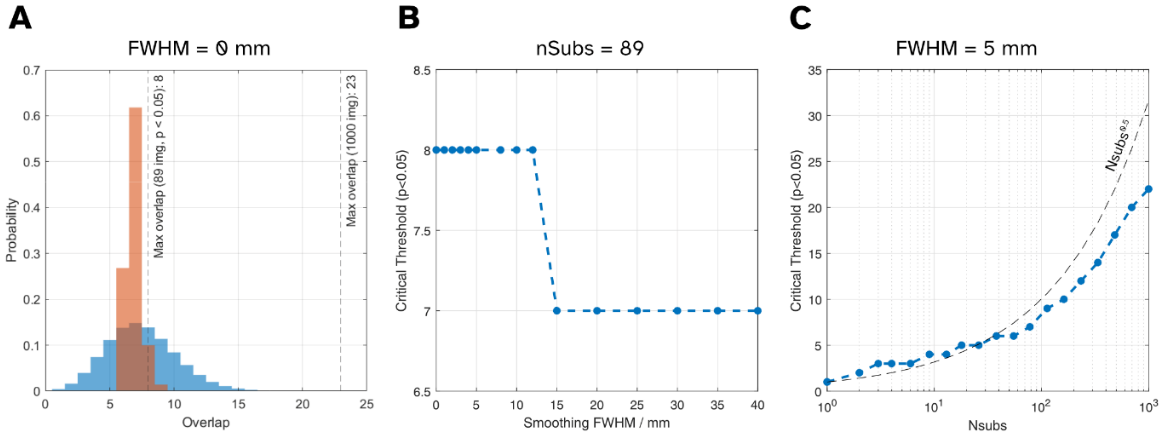


*Figure S1: Results from the empirical null testing which investigated how many subjects overlapping in peak power can be explained from random fields.*

### 2) *Relationship between electrooculography and the brain (continued from the main text)*

Figure S2 shows the relationships between the EOG (and ECG) and the brain. Figure S2A shows the subject-averaged coherence images (as calculated using the DICS method; Gross et al., 2001) for the 200–300 Hz and 300–400 Hz bands. We see that the topography for the HEOG does not change over the bands, but interestingly for the VEOG it moves from the frontal lobe to the pars triangularis and anterior temporal lobes. But again, the coherence is several factors lower than for the HEOG. Figure S2B shows the relationships between the EOG variance in a given band and the beamformer power in the anterior temporal lobes before and after projecting out the EOG signals. The link between the HEOG and source power is again able to become statistically insignificant after projection of the data at the 200–300 Hz band (from $p = 0.046$ to $p = 0.073$). However, we see that in the 300–400 Hz the relationship remains strong after projection (from $p < 0.001$ to $p = 0.001$). A curious observation is that for the VEOG (which showed no relationship in the 100–200 Hz band) now shows significant relationships at 200 Hz and higher. Furthermore, projection out of the data has no significant effect on the relationship (other than a slight reduction). These seems to suggest that generating a sensor-level regression map for SSP on the 100-400 Hz data is biased towards the lower end of the spectrum and not entirely representative of the higher frequencies, something we discuss in the main manuscript regarding independent component analysis. Finally, for completeness, we look at the coherence images between the ECG and the brain in Figure S2C. The coherence scores are very low (less than a part in 1000) and peaks are localised in the cerebellum.

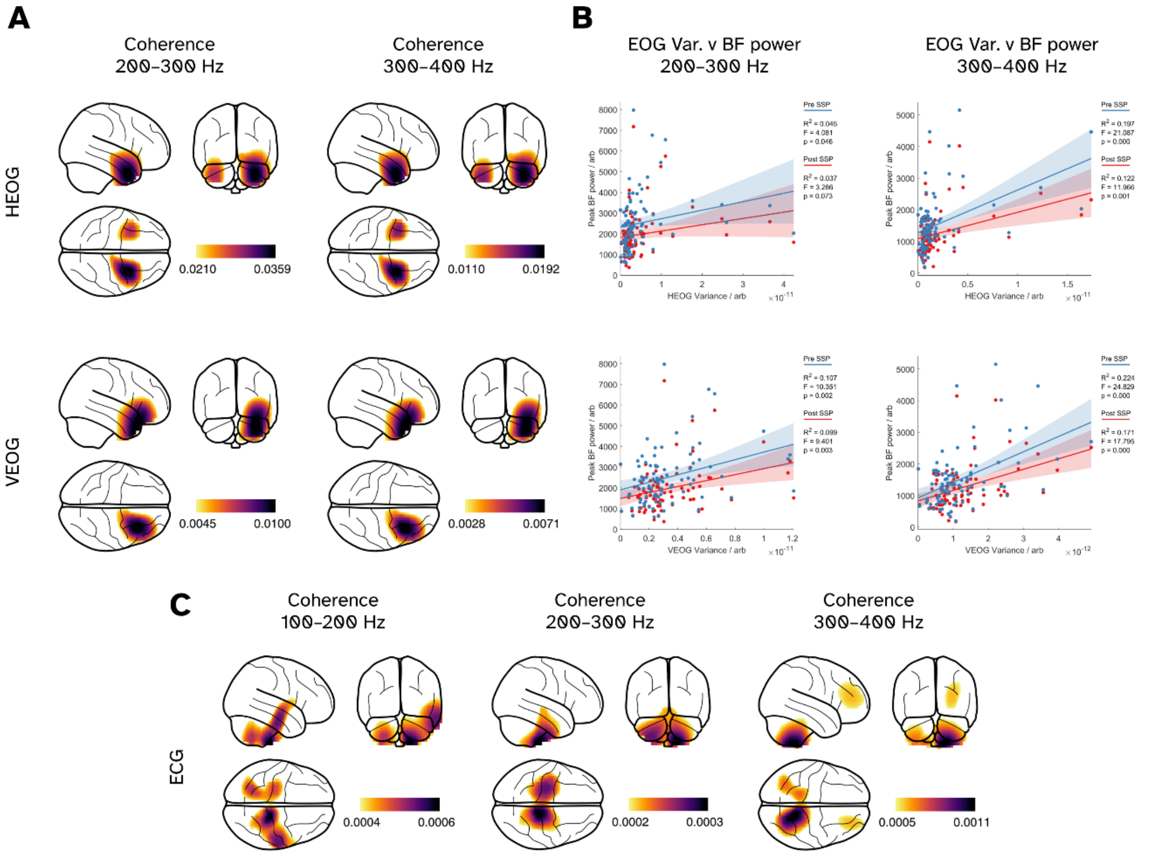


*Figure S2: Additional results showing the relationship between the electrooculogram, electrocardiogram and neural recordings. (A) Coherence between HEOG (top row) and neural data, bottom row represents the coherence between the VEOG and neural data. (B). Linear relationships between the EOG variance (HEOG top, VEOG bottom) for the 200–300 Hz band (left column) and 300–400 Hz band (right column). (C) Coherence between the ECG and the neural data.*

***3) Replication of effects from scene imagination dataset (continued from the main text)***

In addition to the results in the main manuscript, we provide below the unthresholded conjunction maps for this study as from both before and after independent component analysis (FastICA method; Hyvarinen, 1999) has been applied to also investigate where the largest changes in power occur between denoising steps.

*Results*

The results of the analysis on the 100-200 Hz band can be found in Figure S3. To probe the effect of denoising on power localisation in the anterior we opted to contrast between before and after ICA. Whilst Figures S3A and S3B show that our largest overlap across subjects appear in the bilateral superior anterior temporal lobes (left hemisphere, [-54, -6, -10] mm; right hemisphere, [60, -10, -8 mm]), the contrast in Figure S3C shows a marked reduction in peak beamformer power in the anterior temporal lobes and temporal poles after ICA has been applied). For reference, for n=22 subjects our empirical null would set our $p < 0.05$ threshold at 5 subjects or greater, but we have not thresholded the images in the figure. Looking at functional connectivity in Figure S3D, we see the network topography is similar to the results from the HCP dataset. The strongest connections occur from the hippocampal and amygdala ROIs towards the temporal poles. We also observe interhemispheric connections between the left amygdala ROI and right parahippocamal area right olfactory areas. We see a similar effect in the 200-300 Hz bands as well (Figure S4).

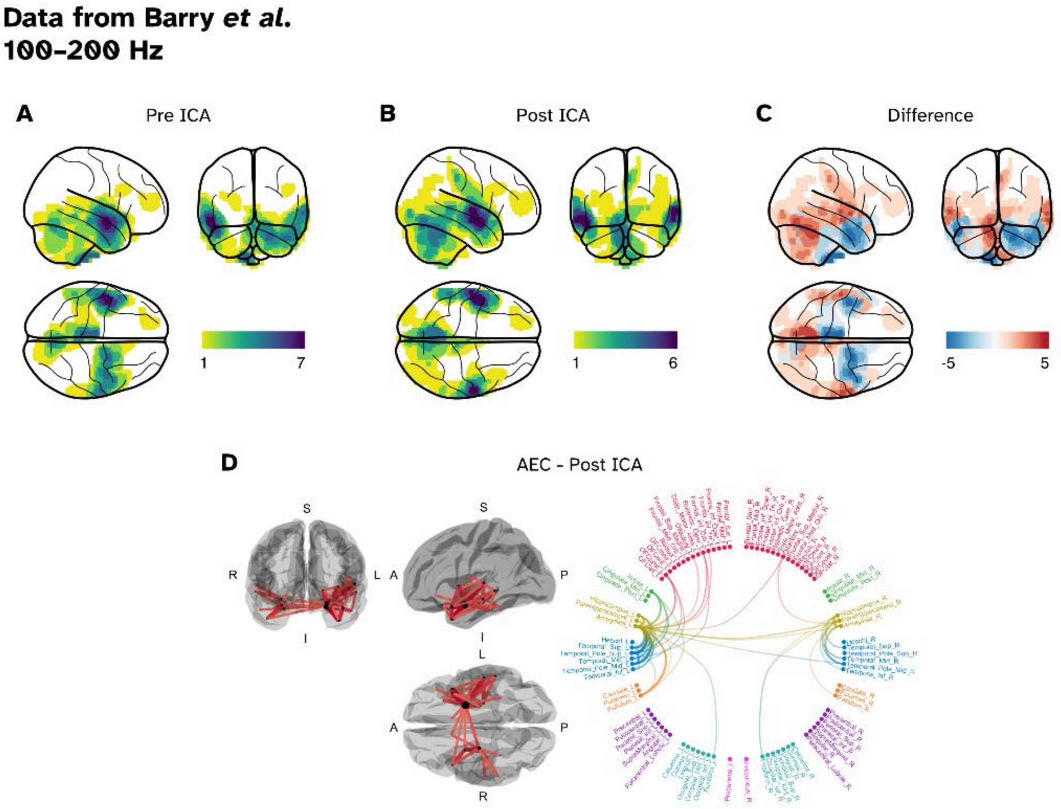


*Figure S3: Replication of the localisation of high frequency power in the 100–200 Hz band in a separate dataset (n=22; Barry et al., 2019). (A) Conjunction of peak power across the subjects prior to denoising via ICA. (B) Conjunction of peak power once ICA has been applied. (C) Glass brains showing the difference in total overlap before and after ICA, showing a reduction within the temporal poles. (D) Subject-averaged Amplitude Envelope Coupling connectome. Looking at the top 50 strongest connections, we observe a similar topography to the networks from the HCP dataset in the main manuscript.*

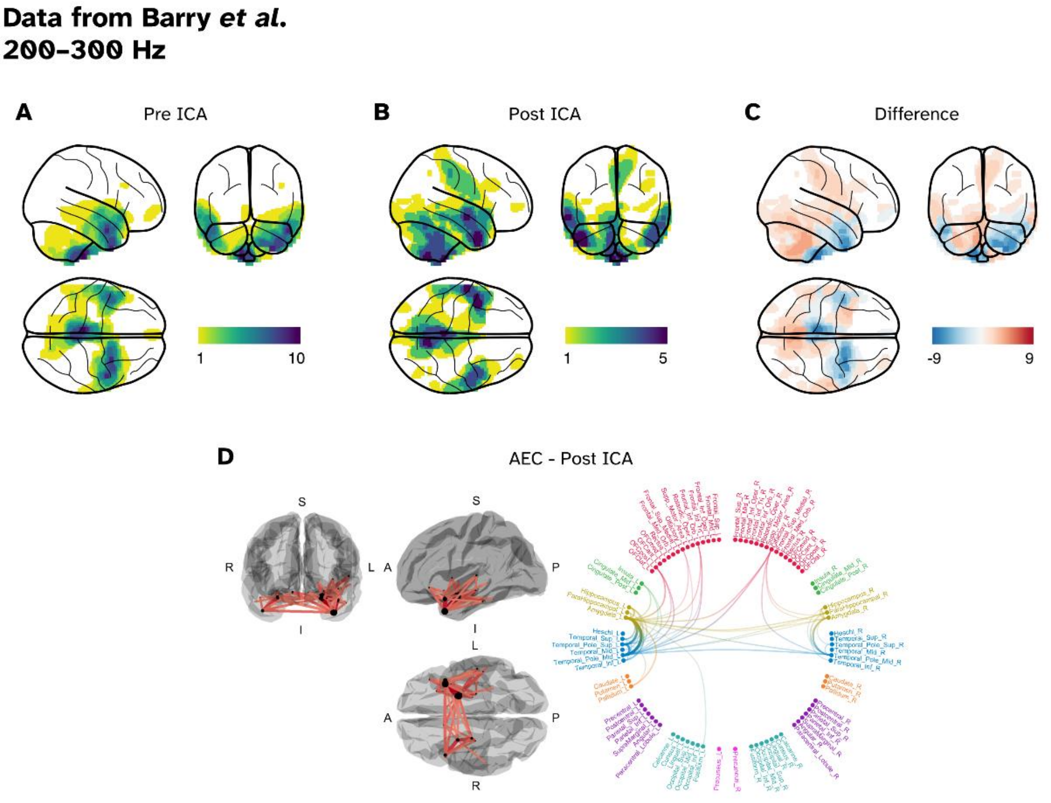


*Figure S4: Replication of the localisation of high frequency power in the 200–300 Hz band in a separate dataset (n=22; Barry et al., 2019). (A) Conjunction of peak power across the subjects prior to denoising via ICA. (B) Conjunction of peak power once ICA has been applied. (C) Glass brains showing the difference in total overlap before and after ICA, showing a reduction within the temporal poles. (D) Subject-averaged Amplitude Envelope Coupling connectome. Looking at the top 50 strongest connections, we observe a similar topography to the networks from the HCP dataset in the main manuscript.*

### *4) The connectome of empty room recordings*

As a control, we projected the empty room recordings (collected prior to a subject entering the MEG) through the beamformer weights of the subjects' resting state recordings and generated a "noise" connectome from the data. The top 50 connections for the three bands are shown in Figure S5. The topographies of the networks do not resemble those seen within the resting state datasets as well as the connectivity being an order of magnitude lower in strength.

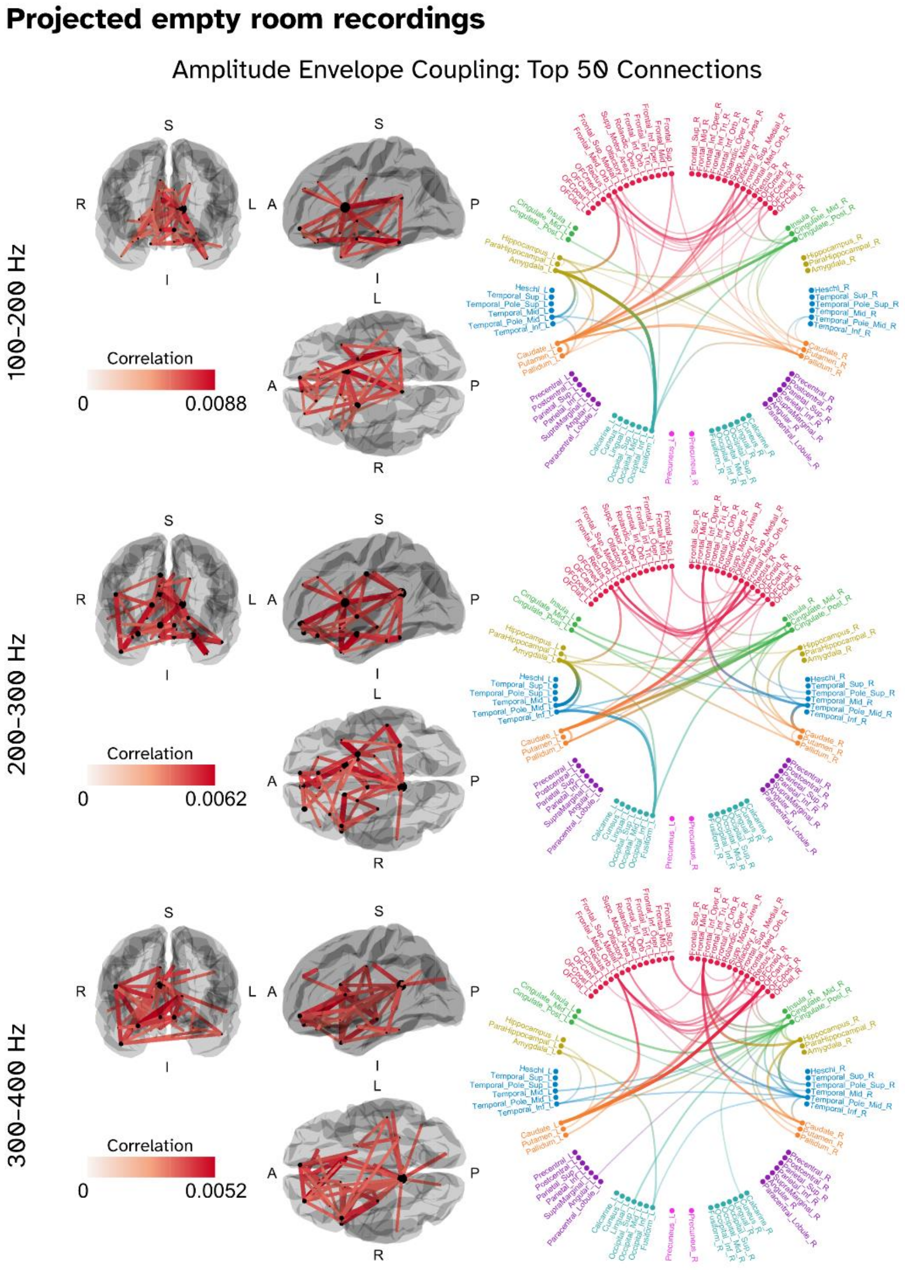


*Figure S5: Connectomes based on the HCP empty room recordings projected through the beamformer weights of the corresponding resting state recordings. The topographies of these networks are distinct to the true resting state networks observed.*
